# Divergent effects of acute and prolonged interleukin 33 exposure on mast cell IgE-mediated functions

**DOI:** 10.1101/463950

**Authors:** Elin Rönnberg, Avan Ghaib, Carlos Ceriol, Mattias Enoksson, Michel Arock, Jesper Säfholm, Maria Ekoff, Gunnar Nilsson

## Abstract

**Background:** Epithelial cytokines, including IL-33 and TSLP, have attracted interest because of their roles in chronic allergic inflammation-related conditions such as asthma. Mast cells are one of the major targets of IL-33, to which they respond by secreting cytokines. Most studies performed thus far have investigated the acute effects of IL-33 on mast cells.

**Objective:** The objective of this study is to investigate how acute versus prolonged exposure of human mast cells to IL-33 and TSLP affects mediator synthesis and IgE-mediated activation.

**Methods:** Human lung mast cells (HLMCs), cord blood-derived mast cells (CBMCs), and the ROSA mast cell line were used for this study. Surface receptor expression and the levels of mediators were measured after treatment with IL-33 and/or TSLP.

**Results:** IL-33 induced the acute release of cytokines. Prolonged exposure to IL-33 increased while TSLP reduced intracellular levels of tryptase. Acute IL-33 treatment strongly potentiated IgE-mediated activation. In contrast, four days of exposure to IL-33 decreased IgE-mediated activation, an effect that was accompanied by a reduction in FcεRI expression.

**Conclusion & Clinical Relevance:** We show that IL-33 plays dual roles for mast cell functions. The acute effect includes cytokine release and the potentiation of IgE-mediated degranulation, whereas prolonged exposure to IL-33 reduces IgE-mediated activation. We conclude that mast cells act quickly in response to the alarmin IL-33 to initiate an acute inflammatory response, whereas extended exposure to IL-33 during prolonged inflammation reduces IgE-mediated responses. This negative feedback effect suggests the presence of a novel IL-33 mediated regulatory pathway that modulates IgE-induced human mast cell responses.

## Introduction

Compelling evidence suggests that epithelial cell-derived cytokines, such as thymic stromal lymphopoietin (TSLP) and interleukin (IL) 33 (IL-33), are strongly involved in the initiation and/or perpetuation of allergy and chronic inflammatory lung diseases such as asthma ^1–3^. Data have accumulated over the years from extensive investigations performed in experimental models, both in vivo and in vitro, as well as from genome wide association studies ^4^ and clinical trials ^5^. Both TSLP and IL-33 are released from epithelial cells in response to pathogens, environmental pollutants and allergens, or, in the case of IL-33, as a result of cell damage. Cells that respond to TSLP and IL-33 include T-lymphocytes, type 2 innate lymphoid cells, eosinophils, basophils and mast cells, all of which are often associated with type 2 immune responses, such as allergies ^6^.

Mast cells are normally located just beneath epithelial cells, and in asthma, they are also found within the intraepithelial cell layer, and they are therefore capable of rapidly responding to TSLP and IL-33 released from epithelial cells ^7^. In contrast to allergen-induced cross-linkage of the IgE receptor, neither TSLP nor IL-33 causes acute mast cell exocytosis, i.e., the release of histamine, proteases and other mediators stored in the granules. Instead, Th2 cytokines are prominently induced in response to TSLP and IL-33 ^8–10^. Interestingly, IL-33 is the sole alarmin that is released from damaged cells that can immediately activate mast cells to induce an inflammatory response and recruit granulocytes ^11,12^. Whether mast cells also synthesize lipid mediators, such as prostaglandins and leukotrienes, in response to TSLP and IL-33 is less clear and might depend on species differences and/or the type of mast cell ^8,10,11,13–16^. Although it does not induce mast cell degranulation on its own, IL-33 increases the synthesis and therefore the amount of pre-stored granule mediators ^17,18^ and augments IgE-mediated mast cell activation ^15,19,20^, and it can therefore potentially aggravate an allergic reaction.

Most of the pioneering experimental studies that have analyzed the effects of TSLP and IL-33 on mast cell degranulation and cytokine production have explored the acute treatment of mast cells, i.e., a timescale of minutes to a few hours ^21^. Those results can probably be accurately transferred to an acute inflammatory situation in which mast cells should respond to “danger”, such as trauma. However, during prolonged chronic inflammation, such as asthma, the expression of IL-33 is increased in both epithelial cells and airway smooth muscle cells ^22,23^, two compartments of the asthmatic lung that are associated with increased mast cell numbers ^24–26^. We therefore asked how human mast cell functions are affected by acute and prolonged exposure to IL-33 and/or TSLP. The expression profiles of receptors for IL-33 and TSLP were analyzed in human lung mast cells (HLMCs), primary developed mast cells, and mast cell lines. The effects of acute or prolonged exposure to IL-33 and/or TSLP on mediators, receptors and IgE-mediated activation were analyzed. Our results reveal that IL-33 increases the FcεRI-mediated response when cells are concomitantly exposed to IL-33 and an antigen, whereas prolonged exposure to IL-33 inhibits FcεRI expression and thereby diminishes IgE-mediated mast cell degranulation. IL-33 may therefore play a significant role in the regulation of mast cell reactivity in IgE-associated chronic inflammation.

## Methods

### Ethical approval

The local ethics committee approved the experiments involving human subjects, i.e., the collection of lung tissue from patients undergoing lobectomies, and all patients provided informed consent. In accordance with Swedish legislation, ethics approval was not needed for the anonymous collection of cord blood because the samples cannot be traced to a specific person.

### Cell culture and in vitro stimulations

The human mast cell line ROSAWT KIT 27 was cultured in IMDM supplemented with 10% fetal calf serum, 2 mM L-glutamine, 100 μg/mL streptomycin, 100 IU/mL penicillin and 80 ng/mL murine stem cell factor (SCF). Cord blood-derived mast cells (CBMCs) were cultured as previously described ^28^. Single cell suspensions obtained from human lung tissue, for the analysis of HLMCs, were obtained as previously described^29^ and maintained in RPMI 1640 medium (Sigma Aldrich) supplemented with 10% fetal calf serum, 100 ng/ml hSCF, 0.01 M HEPES, 0.5x nonessential amino acids, 2 mM L-glutamine, 100 units/ml penicillin, 0.1 mg/ml streptomycin and 48 μM β-mercaptoethanol (Sigma Aldrich). Cells were stimulated with 10 ng/ml TSLP and/or 10 ng/ml IL-33 (Peprotech, Rocky Hill, NJ, USA). The cytokine concentration was chosen based on published data^8,10,13^. The response was analyzed after either 1 hour of stimulation or 4 days with daily addition of the cytokines without media change. To measure the levels of FcεRI receptor (in ROSA cells and CBMCs) and the amount of degranulation induced by FcεRI crosslinking (CBMCs), 10 ng/ml IL-4 (Peprotech) was added 4 days prior (unless otherwise stated) and 1 µg/ml human IgE (Calbiochem, Minneapolis, MN, USA) was added one day prior to crosslinking. Cells were crosslinked with various concentrations of anti-IgE antibody (Sigma), and calcium ionophore A23187 (2 µM, Sigma) was used as a positive control for activation. In some experiments (indicated in the figure legends) performed to measure lipid mediators, the cells were pretreated with 10 ng/ml IL-4 and 5 ng/ml IL-3 for 4 days.

### Measurement of mediator release

Released histamine was measured using a histamine release test kit according to the manufacturer’s instructions (RefLab, Copenhagen, Denmark). Briefly, this test is based on the adsorption of histamine to glass fiber-coated microtiter plates. The glass fibers bind histamine with high affinity and selectivity. The plates were sent to RefLab, and histamine was released and detected fluorometrically (OPA-method) by HISTAREADER™ 501-1. PGD_2_ was measured using a Prostaglandin D_2_-MOX ELISA kit (Caymen Chemical, Ann Arbor, MI. USA), and the levels of IL-1β, IL-5, MCP-1, MIP-1α, GM-CSF and TNF were analyzed with Luminex (BioRad, Hercules, CA, USA).

### Flow cytometry

The following antibodies were used for surface staining: ST2-FITC (clone B4E6, MD Bioproducts, Zürich, Switzerland), IL7R-PE (clone A019D5, Biolegend, San Diego, CA, USA), TSLP-R-PE (clone 1B4, Biolegend), FcεRIα-PE (clone AER-37 (CRA-1), Biolegend) and CD63-Pe-Cy7 (Clone H5C6, BD Biosciences, San Jose, CA, USA). Human lung cells were stained with BD Horizon™ Fixable Viability Stain 450 (BD Biosciences) and CD45-V500 (Clone HI30, BD Biosciences), CD14-APC-Cy7 (Clone M5E2, Biolegend), and CD117-APC (clone 104D2, BD Biosciences) antibodies; and mast cells were gated as live, CD45^+^, CD14^low^ CD117^high^. For intracellular staining, cells were fixed with 4% PFA and permeabilized using PBS-S buffer (0.1% saponin in PBS with 0.01 M HEPES). Unspecific binding was blocked using blocking buffer (PBS-S with 5% dry milk and 2% FCS). The cells were thereafter stained with tryptase antibodies (clone G3, Millipore, Burlington, MA, USA) conjugated in-house with an Alexa Flour 647 Monoclonal antibody labeling kit (Invitrogen), chymase antibodies (clone B7, Millipore) conjugated in-house with a PE Conjugation Kit (Abcam, Cambridge, UK) or CPA3 antibodies (clone CA5, a kind gift from Andrew Walls, Southampton, UK) conjugated in-house with an Alexa Fluor™ 488 Antibody Labeling Kit (Thermo Fisher Scientific, Waltham, MA, USA). The cells were analyzed using a BD FACSCanto system (BD, Franklin Lakes, NJ, USA), and FlowJo software (FlowJo LLC, Ashland, OR, USA) was used for flow cytometry data analysis.

### Quantitative PCR

RNA was extracted using an RNeasy Plus Mini Kit (Qiagen, Hilden, Germany), cDNA was prepared using an iScript cDNA synthesis kit (Bio-Rad), and qPCR was performed using iTaq Universal SYBR Green Supermix (Biorad) on a CFX96 Real-time system (Biorad). The following primers were used: GAPDH (5'-CCACATCGCTCAGACACCAT-3' and 5'-GGCAACAATATCCACTTTACCAGAG-3'), FcεRIα(5'-CGCGTGAGAAGTACTGGCTA-3' and 5'-TGTGACCTGCTGCTGAGTTG-3'), FcεRIβ(5'-TGCAGTAAGAGGAAATCCACCA-3' and 5'-TGTGTTACCCCCAGGAACTC-3'), and FcεRIγ(5'-CCAGCAGTGGTCTTGCTCTT-3' and 5'-AGGCCCGTGTAAACACCATC-3'). The results were calculated using the ΔΔCT method, and CT values were normalized to the housekeeping gene GAPDH and related to the unstimulated control.

### Statistical analysis

Data are shown as the mean ± the standard error of the mean (SEM). Statistical analyses were performed with GraphPad Prism software version 7.0b. When comparing 2 groups, Student’s *t* test was performed, and when more than 2 groups were compared, one-way or two-way ANOVA with Bonferroni’s post hoc test was performed (*, P < 0.05; **, P < 0.01; ***, P < 0.001; ****, P < 0.0001).

## Results

### Expression of receptors for IL-33 and TSLP on mast cells

The expression levels of different receptors for IL-33 (ST2) and TSLP (TSLP-R and IL7R) were analyzed in mast cell line ROSA and primary CBMCs and HLMCs by flow cytometry. The expression level of TSLP-R was higher in ROSA cells than in the primary CBMCs and HLMCs (Fig 1A). IL7R staining was very low in all cells analyzed (Fig 1B) but all cells had detectable levels of the IL-33 receptor ST2 (Fig 1C). When the mast cells were treated for 4 days with IL-33, the expression level of TSLP-R increased, and this effect was counteracted by TSLP, possibly via the internalization of the receptor (Fig 1D). The addition of TSLP also reduced surface staining for IL7R (Fig 1E). None of the cytokines had any significant effect on ST2 receptor expression (Fig 1F).

**Figure 1.**
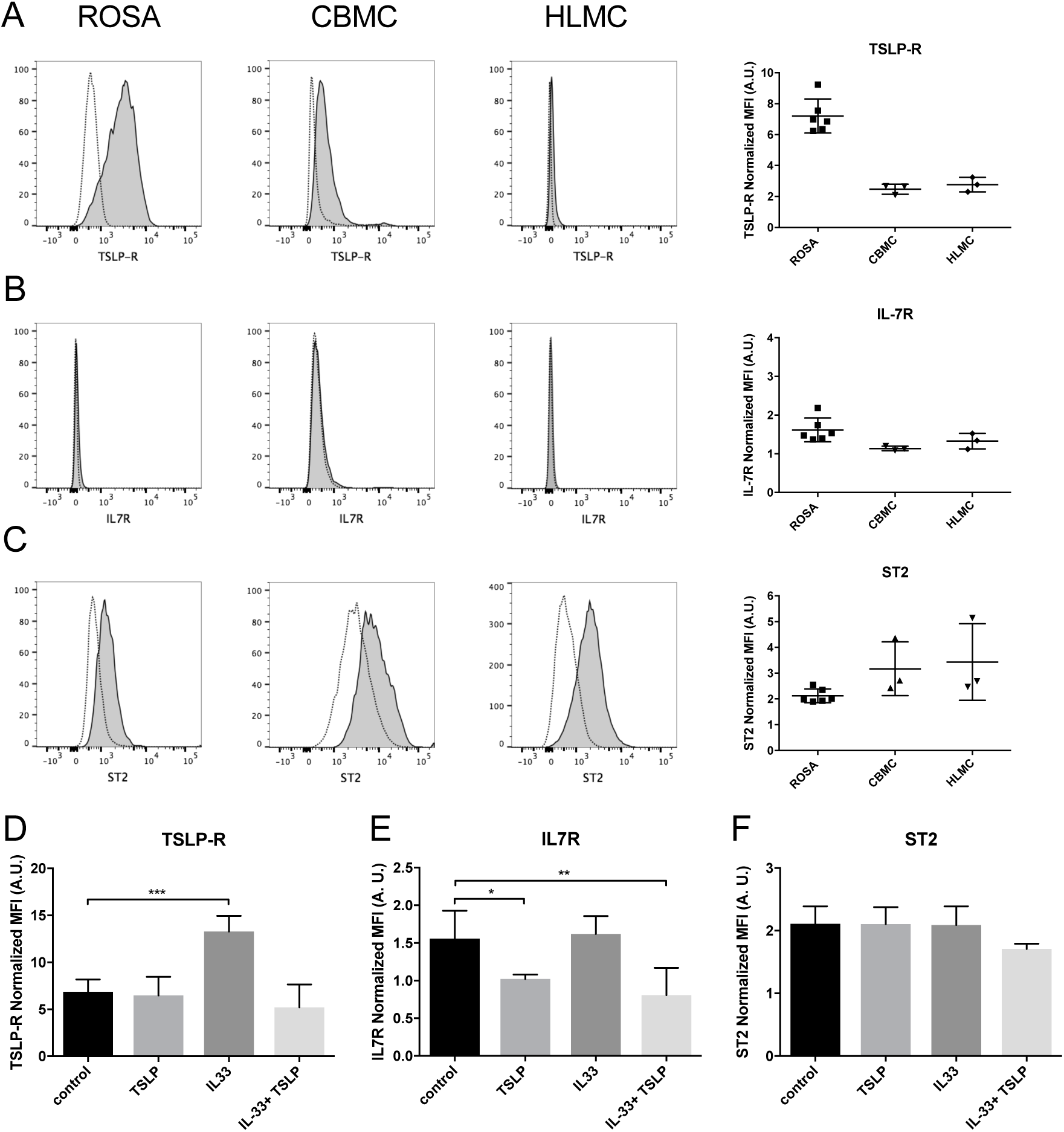
Expression of receptors for IL-33 and TSLP on various human mast cells. Surface expression of TSLP-R (A), IL7R (B) and ST2 (C) on ROSA, CBMC and HLMCs was measured by flow cytometry. Representative histograms in which dotted lines represent the respective isotype controls are shown for different cell lines to the left, and quantification of the data is shown to the right (A-C). ROSA cells were treated with 10 ng/ml IL-33, TSLP or a combination or both repeatedly for four days; thereafter, surface expression of TSLP-R (D), IL7R (E) and ST2 (F) was measured by flow cytometry. The median fluorescent intensity (MFI) of the receptors was normalized to the respective isotype control. Data shown were pooled from 3 independent experiments, n=3-6.

### Degranulation after prolonged exposure to IL-33 and TSLP

It has been reported that neither IL-33 nor TSLP causes mast cell degranulation ^8–10,19,30^, and our results confirm this finding because no histamine was released after 1 hour of stimulation (Fig 2A, C); however, after 4 days of stimulation, IL-33 caused a small but significant increase in the release of histamine in CBMCs (Fig 2D), and the combined addition of IL-33 and TSLP induced histamine release in ROSA cells (Fig 2B).

**Figure 2.**
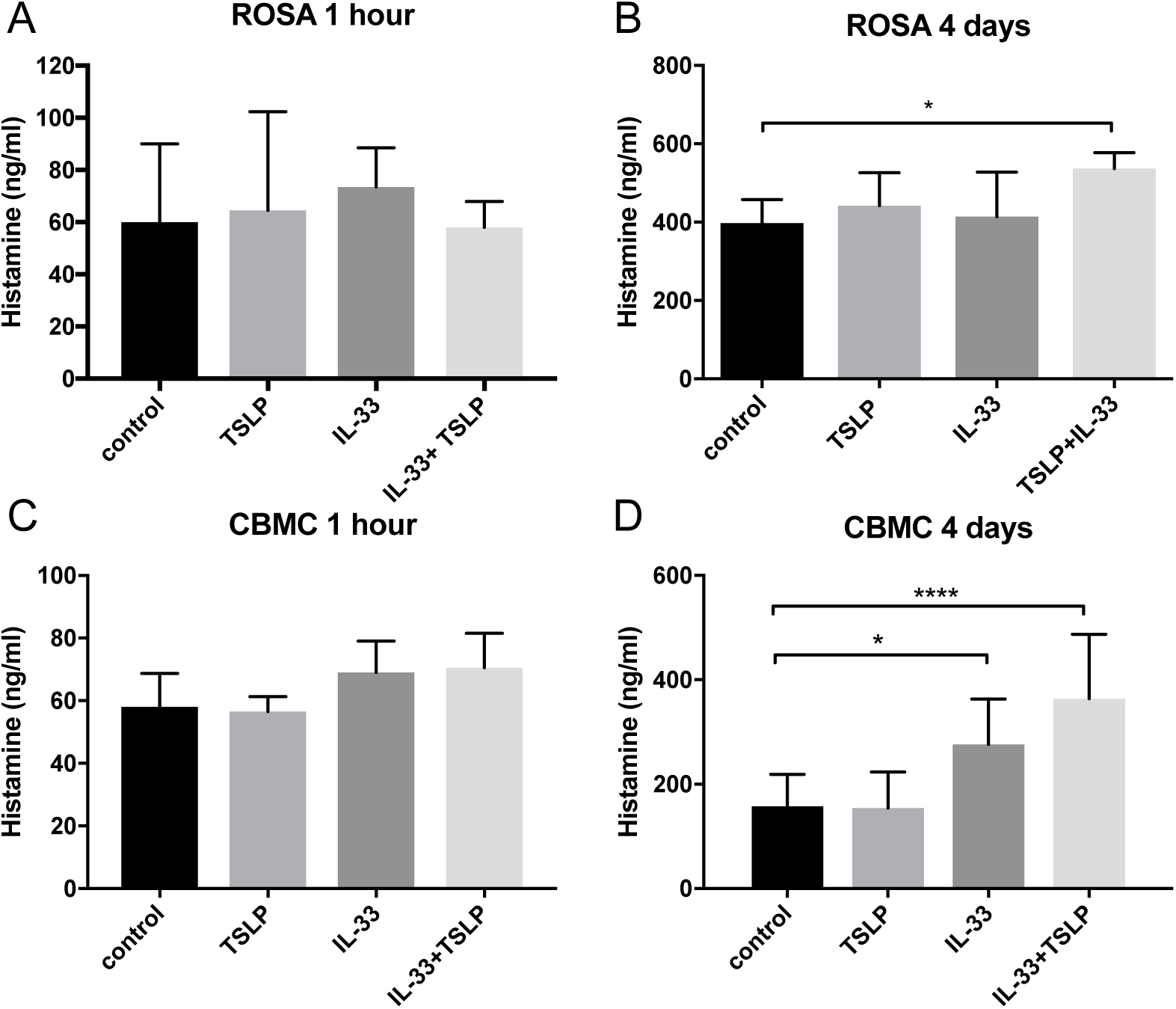
Histamine release induced by prolonged exposure to IL-33. ROSA cells (A-B) or CBMCs (C-D) were treated with 10 ng/ml IL-33, TSLP or a combination of both for 1 hour (A, C) or treated repeatedly (every day) for 4 days (B, D). Thereafter, supernatants were collected, and histamine was measured. Data shown were pooled from 3-5 independent experiments, n=6-10.

### Prostaglandin D_2_ and cytokine release induced by IL-33

The results of reports regarding whether IL-33 induces the release of lipid mediators in mast cells have been inconsistent ^10,11,13–16^. We observed considerable variation in the basal levels of PGD_2_ released from different CBMC cultures: in some, there appeared to be an increase following IL-33 stimulation, whereas there was no change in others. Collectively, our results indicated there was no significant increase overall. Pretreating the cells with IL-4 and IL-3 for 4 days increased the amount of PGD_2_ released but did not increase the number of CBMC cultures that responded to IL-33 (Fig 3B). TSLP did not induce any PGD_2_ release (data not shown). All cultures treated with calcium ionophore A23187 (used as a positive control) exhibited a substantial increase in the amount of PGD_2_ released following this treatment (Fig 3C). There was no detectable release of cysteine leukotrienes in response to IL-33 or TSLP (data not shown). IL-33 caused a significant increase in the cytokines IL-5, GM-CSF and TNF (Fig 3D-F) but did not affect the amount of IL-1β, MCP-1 and MIP-1α released (Fig 3G-I).

**Figure 3.**
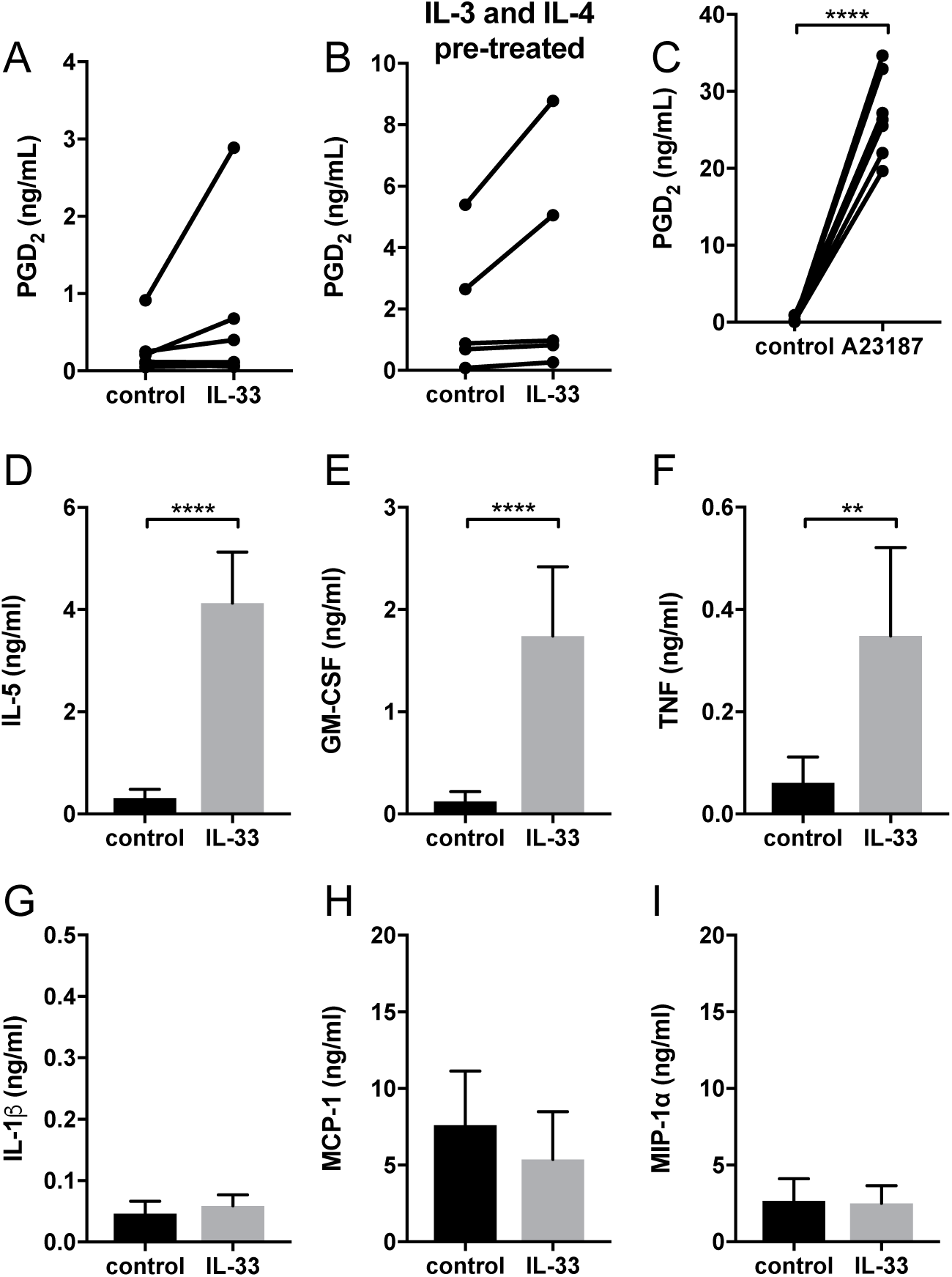
Prostaglandin D_2_ and cytokine release in response to IL-33. CBMCs were treated for 1 hour (A-C) or 24 hours (D-I) with 10 ng/ml IL-33 or 2 μM A23187, and supernatants were collected. PGD_2_ (A-C) was measured using ELISA, and cytokines were measured using Luminex assays (D-I). The CBMCs shown in B were pretreated with 10 ng/ml IL-4 and 5 ng/ml IL-3 for 4 days. Data shown were pooled from 5-9 independent experiments, n=5-9

### Effects on mediator storage by IL-33 and TSLP

Next, we investigated whether the storage of mast cell mediators would be affected by 4 days of exposure to IL-33 and TSLP. Intracellular tryptase was increased by IL-33 but decreased by TSLP in ROSA cells (Fig 4A). IL-33 also increased tryptase storage in CBMCs, while TSLP alone had no effect in these mast cells, but when added with IL-33, it lowered the IL-33-induced increase (Fig 4B). Neither IL-33 nor TSLP had any effect on the storage of chymase, CPA3 or histamine (Fig 4 C-F).

**Figure 4.**
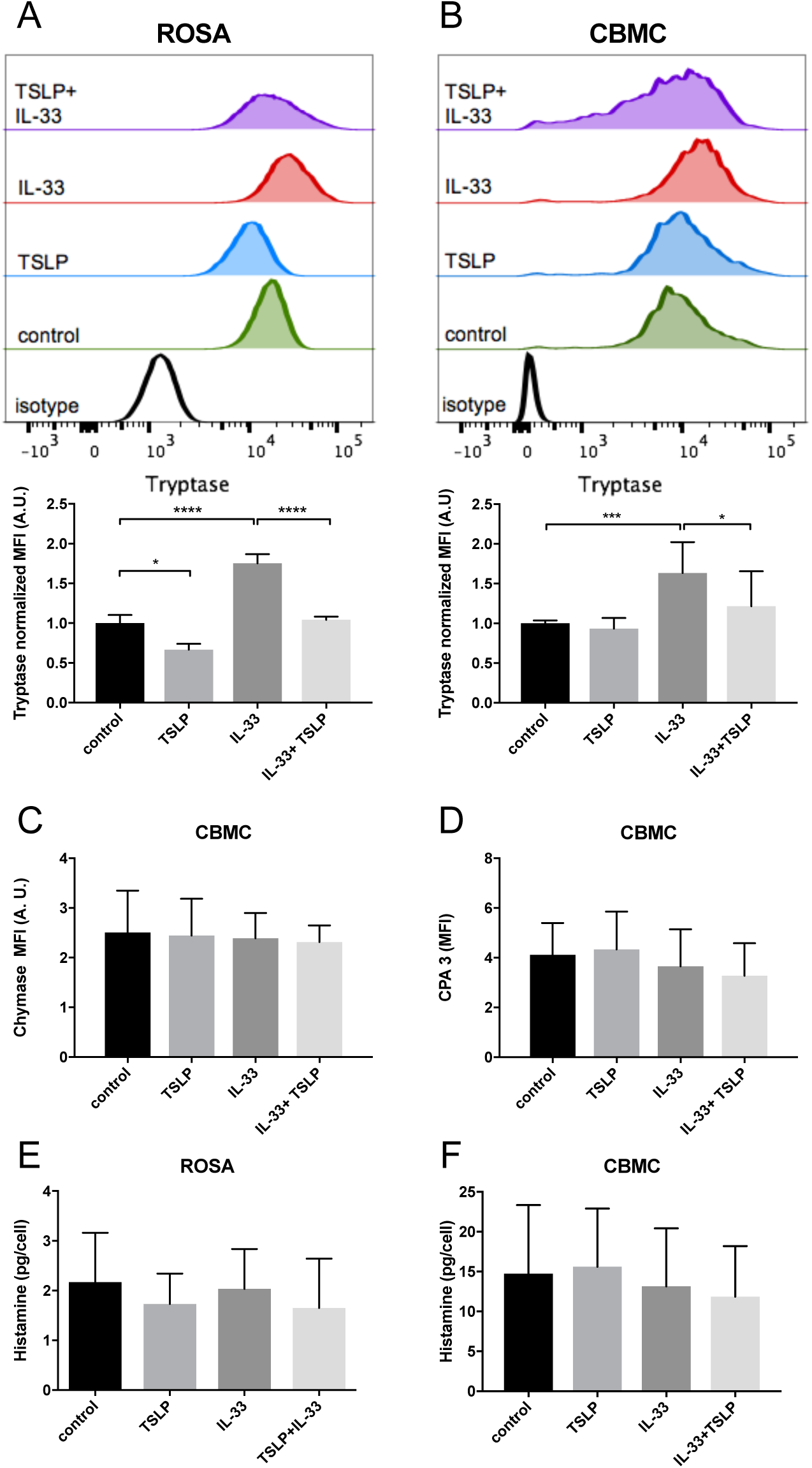
Changes in mediator storage after 4 days of treatment with IL-33 and TSLP. ROSA cells (A, E) or CBMCs (B-D, F) were repeatedly treated every day for four days with 10 ng/ml IL-33, TSLP or a combination of both; thereafter, the levels of the proteases tryptase (A-B), chymase (C) and CPA3 (D) were measured by intracellular flow cytometry staining, and total histamine content (E-F) was measured by a histamine release test kit. The MFI of protease expression was normalized to the respective isotype control. A, a representative of 3 independent experiments, n=3. Data shown were pooled from 5 (B), 2 (C-D), 3 (E) and 6 (F) independent experiments, n=4-12.

### Effects on IgE-mediated degranulation by IL-33

It has previously been demonstrated that IL-33 potentiates IgE-mediated degranulation ^15,17,19,20^. Pretreatment with IL-33 for 1 hour prior to FcεRI crosslinking increased both surface CD63 expression and histamine release (Fig 5 A, C). In contrast, when CBMCs were repeatedly treated with IL-33 for 4 days, IgE-mediated degranulation was significantly decreased, with reduced induction of CD63 surface expression and histamine release compared to the untreated group (Fig 5B, D). Pretreatment with TSLP for 4 days had no effect on IgE-mediated degranulation (data not shown).

**Figure 5.**
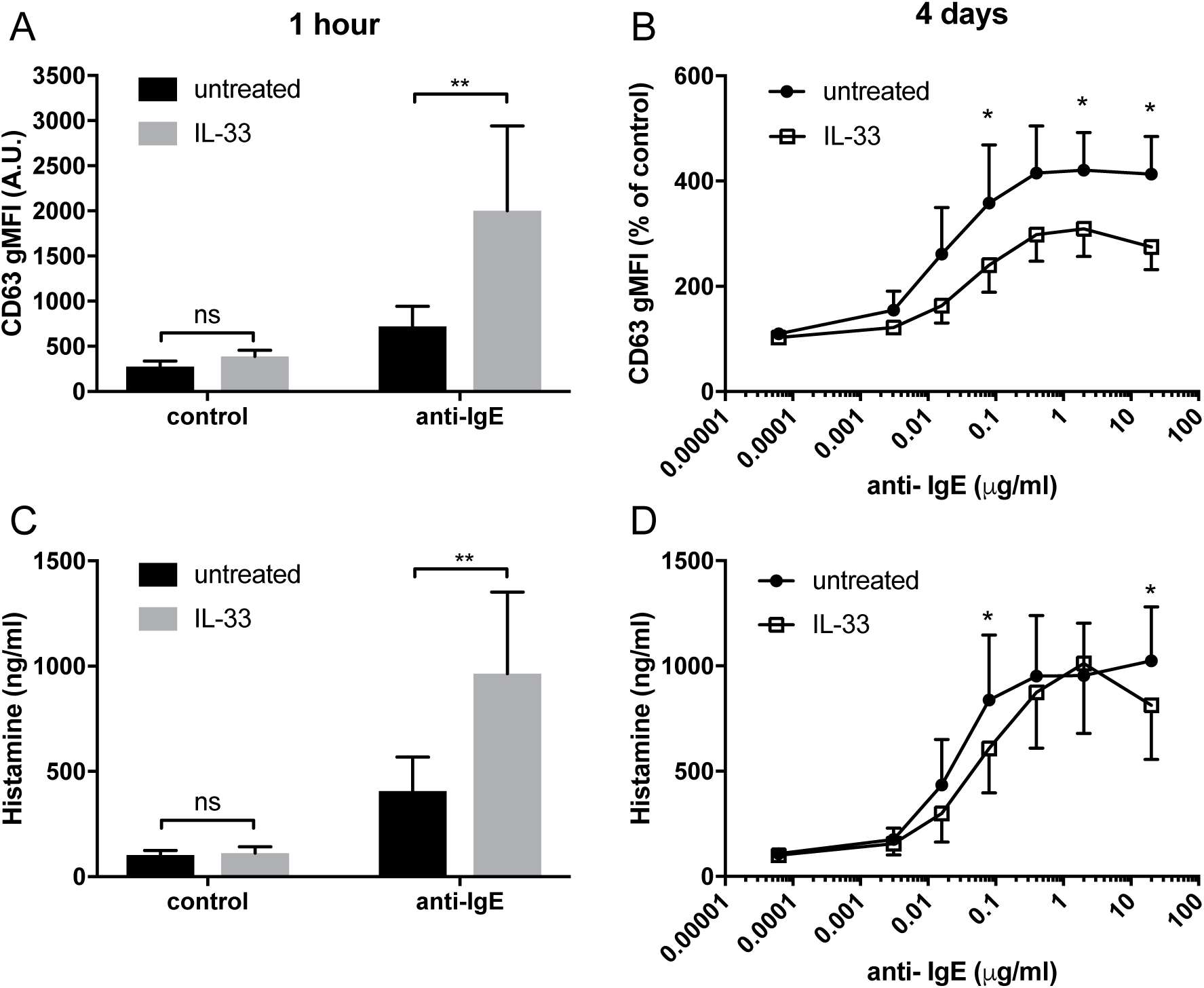
Effect of IL-33 on mast cell degranulation by FcεRI crosslinking. CBMCs were treated with 10 ng/ml IL-33 for 1 hour or repeatedly every day for four days; thereafter, they were stimulated with 0.4 μg/ml (A, C) or various concentrations (B, D) of anti-IgE. Degranulation was measured by the detection of surface CD63 with flow cytometry (A-B) or as histamine release (C-D). Data shown were pooled from 3 independent experiments, n=3-6.

### FcεRI surface receptor expression is decreased by 4 days of IL-33 treatment

To further investigate a possible mechanism that could underlie the observed decrease in degranulation, we next investigated the effect of IL-33 and TSLP on surface expression of the FcεRI receptor. We found that four days of treatment with IL-33 caused a dramatic decrease in the surface expression of the FcεRI receptor in ROSA cells, CBMCs and HLMCs (Fig 6A-F). In ROSA cells, TSLP also caused a significant drop in FcεRI surface staining (Fig 5A, D). Since human mast cells grown in culture express very low levels of FcεRI, we added IL-4 to upregulate the receptor in ROSA cells and CBMCs. To investigate whether IL-33 reduces receptor expression simply by blocking IL-4-mediated receptor upregulation, IL-4 was added four days prior to a media change, and IL-33 and TSLP addition. Also, in this experiment, IL-33 downregulated the FcεRI receptor, indicating that a blockade of IL-4-mediated upregulation is not the mechanism that reduces FcεRI receptor surface expression. However, the reduction of FcεRI by TSLP was absent in this case, indicating that TSLP is blocking the IL-4 mediated upregulation (Fig 6 G-H).

**Figure 6.**
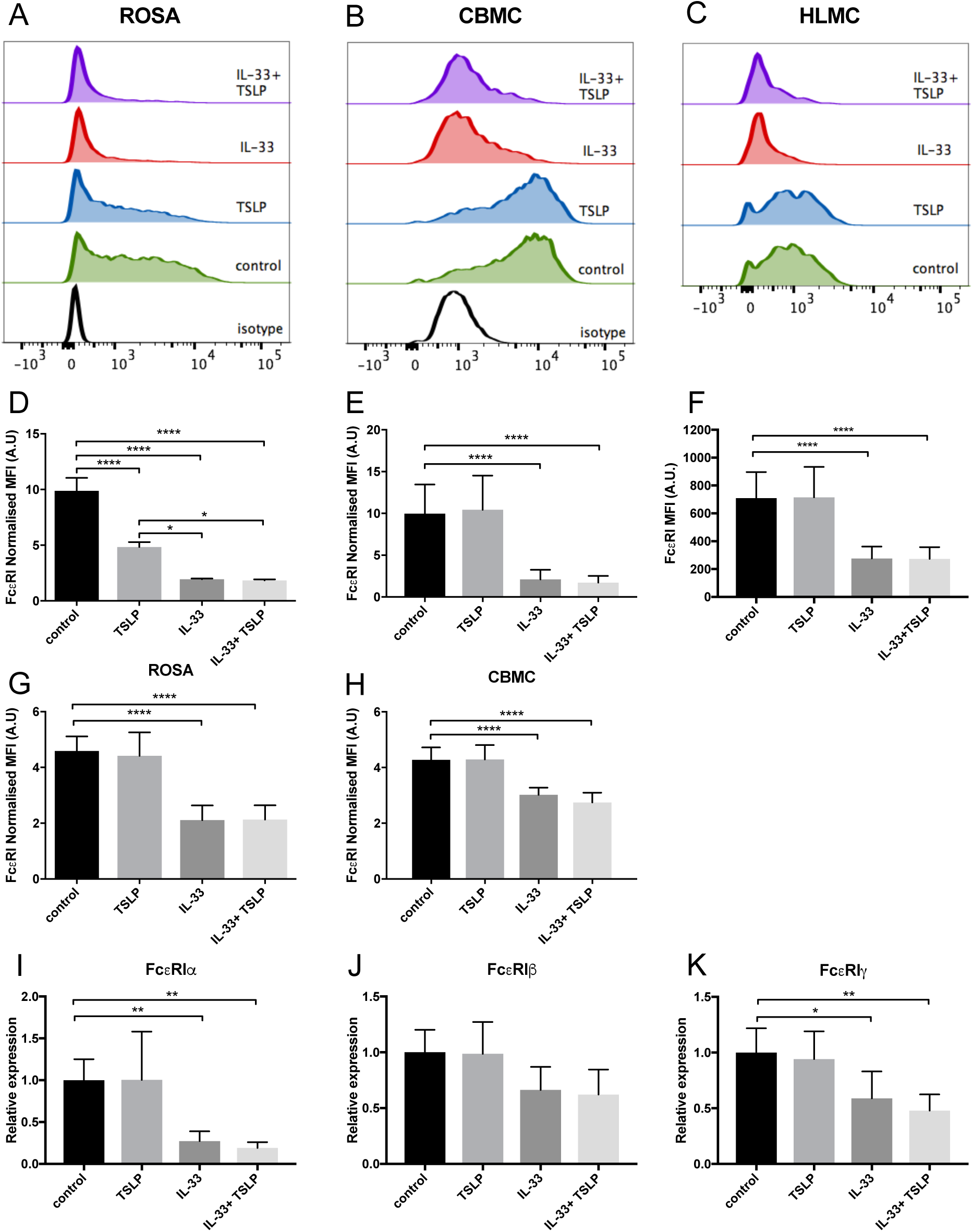
Decrease in FcεRI surface expression and mRNA level by IL-33. ROSA cells (A, D, G), CBMCs (B, E, H, I, J, K) or HLMCs (C, F) were repeatedly treated every day for four days with 10 ng/ml IL-33, TSLP or a combination of both. In addition, IL-4 was added on day 0 (A, B, D, E) or 4 days prior to the addition of IL-33 and TSLP (G-H) to upregulate the FcεRI receptor. Surface FcεRI receptor expression was measured by flow cytometry, representative histograms are shown in A-C and quantification of the expression is shown in D-F. The MFI of FcεRI was normalized to that of the isotype control (D-E, G-H). The mRNA expression level of the different subunits of the FcεRI receptor were quantified using qPCR (I-K). Expression was normalized to that of the housekeeping gene GAPDH and thereafter to the unstimulated control. Data shown were pooled from 3-6 independent experiments, n=6-10.

### IL-33 decreases the expression of *FcεRI*-*α* and -*γ* subunits

The FcεRI receptor consists of four chains, including one α-, one β-and two *γ*-chains, which are regulated at both the protein and mRNA levels ^31^. We therefore next investigated whether IL-33 affects the mRNA expression of the different subunits of this receptor. We found that four days of treatment with IL-33 significantly reduced the levels of the FcεRI-α and -γ subunits, indicating that the relevant regulatory mechanism occurs at the mRNA level (Fig 6 I-K).

## Discussion

Compelling evidence indicates that IL-33 acts as an alarmin to activate mast cells in an acute manner, resulting in the release of pro-inflammatory mediators ^32^. In the present study, we demonstrate an opposite effect of IL-33 by which prolonged exposure to IL-33, down-regulates the high-affinity IgE receptor and reduces the allergic response (Fig 5, 6). Thus, IL-33 appears to have divergent functions on mast cells, including an acute effect by which it induces the release of cytokines (Fig 3) and consequentially acute inflammation and a dampening effect observed in cells exposed to IL-33 for a longer period, such as during chronic inflammation^1^.

We also investigated the surface expression of receptors for IL-33 and TSLP, including ST2, TSLP-R and IL7R, in the mast cell line ROSA as well as primary CBMCs and HLMCs. Similar to previously published reports, we found that all of these cells expressed the ST2 receptor (Fig 1C) ^8,10,14,33^; however, contrary to Kaur et al.^33^, we did not observe any increase in the surface expression of ST2 after IL-33 treatment (Fig 1F), possibly because we exposed the cells to IL-33 for different periods of time (24 hours versus 4 days). TSLP-R surface staining was stronger in the mast cell lines than in primary mast cells (Fig 1A), and the ROSA mast cells were the only cells that responded to TSLP, with no response observed in the primary cells (Fig 4A-B, Fig 6A-F). However, exposure to IL-33 upregulated TSLP-R surface expression (Fig 1D), and in some cases, we also observed that TSLP affected primary cells when added in combination with IL-33 (Fig 2D, Fig 4B), possibly because of the upregulation of its receptor. IL7R surface staining was very low in all cells, but when cells were treated with TSLP, it was even lower, suggesting that although it is expressed at very low levels, it is still functional and internalized upon ligand binding (Fig 1E).

Several studies have previously demonstrated that IL-33 induces the release of cytokines from mast cells ^8,10,11,14,15,19,20,30,34^. Studies have also shown that IL-33 induces the release of lipid mediators ^11,13,14,16^, while other studies have reported no lipid mediator release in response to IL-33 ^8,10,15^. We also observed that IL-33 induced the release of cytokines (Fig 3D-I); however, there was high donor to donor variation in PGD_2_ release, resulting in no significant change overall (Fig 3A-B). The reason for this variation between donors remains unresolved, and further studies are needed to determine why some patients do not release PGD_2_ in response to IL-33.

IL-33 has been proposed to be important for the maturation of mast cells because it increases storage of the mast cell proteases tryptase and CPA3 as well as the amine serotonin ^10,16–18^. In our experiments, 4 days of exposure to IL-33 also increased, whereas TSLP decreased, the intracellular level of tryptase. These findings are in contrast to the results presented by Lai et al., who found that storage of tryptase was increased by TSLP ^16^. This difference could be because we used mature CBMCs (more than 8 weeks in culture), while they added TSLP when the cultures of cord blood cells were begun in order to develop them into mast cells. They also cultured the cells in the presence of TSLP for 3 weeks, while we did so for only 4 days. Interestingly, this was the only experiment in our study in which IL-33 and TSLP exerted opposing effects. We did not observe any change in chymase and CPA3 expression after 4 days of stimulation, in agreement with the results presented in Lai et al^16^. However, they reported that CPA3 expression was increased after 3 weeks of treatment. Human mast cells do not contain as much serotonin as is contained in mouse mast cells, but they store more histamine ^35^. We therefore investigated whether the storage of histamine was affected by IL-33 and TSLP, but we observed no change after 4 days of treatment (Fig 4E-F).

Previous studies have shown that neither IL-33 nor TSLP alone induces acute mast cell degranulation^20,36,37^, and this finding was also confirmed in this study (Fig 2A, C). However, 4 days of exposure to IL-33 induced partial exocytosis and histamine release (Fig 2B, D). Similar to previous studies ^20,37^, we found that 1 hour of pretreatment with IL-33 strongly potentiated the mast cell degranulation induced by crosslinkage of the IgE-receptor (Fig 5A, C). In contrast, 4 days of exposure to IL-33 had the opposite effect, with the treated cells exhibiting less degranulation than was observed in the untreated group (Fig 5B, D). Jung et al. also showed that prolonged exposure to IL-33 induced a hyporesponsive phenotype in mast cells. They investigated the cause of this hyporesponsiveness in mouse mast cells and observed that while there was no difference in FceRI receptor expression, but Hck expression was decreased ^38^. We did not observe any change in Hck expression (data not shown) in human mast cells, but FceRI surface receptor expression dropped dramatically in both human ROSA cells and primary CBMCs as well as in primary mast cells isolated from human lungs (Fig 6A-F). FceRI expression can be regulated at both the mRNA and protein level ^31^; IL-33 mediated the downregulation of FceRI by decreasing the mRNA expression levels of its -α and -γ subunits (Fig 6I-K). Since IL-33 activates mast cells to release various cytokines it is not clear if the observed long-term effect of IL-33 is a direct effect or if it is a secondary consequence of the mediators that are released by IL-33 activation. This warrents further investigation to clarify.

Mast cells are very potent pro-inflammatory cells, and the systemic activation of mast cells can lead to anaphylaxis and potentially death. Mast cell activation must therefore be tightly regulated ^39^. IL-33 is a potent pro-inflammatory cytokine, and it is clear that acute exposure to IL-33 activates mast cells and increases their responsiveness to antigens. This is a potentially dangerous situation, and we therefore suggest that after prolonged exposure to IL-33, mast cells down-regulate the FceRI receptor in a negative feedback loop to prevent damage caused by excessive mast cell activation.

## Acknowledgement

We thank SOBI, Stockholm, Sweden, for generously gifting SCF and Andrew Walls for generously gifting the CPA3 antibody. This study was supported by grants from the Swedish Research Council; the Heart-Lung Foundation; the Ollie and Elof Ericssons foundation; the Ellen, Walter and Lennart Hesselman’s foundation; Tore Nilssons Foundation; the Lars Hiertas memory fund; the Konsul Th C Burghs Foundation; the Tornspiran Foundation; the O. E. and Edla Johanssons Foundation; the Swedish Society for Medical Research; The ChAMP (Centre for Allergy Research Highlights Asthma Markers of Phenotype) consortium funded by the Swedish Foundation for Strategic Research; the AstraZeneca & Science for Life Laboratory Joint Research Collaboration; and the Karolinska Institutet.

## Conflict of interest statement

The authors declare that this research was conducted in the absence of any commercial or financial relationships that could be construed as a potential conflict of interest.

